# Variational Autoencoder-based Model Improves Polygenic Prediction in Blood Cell Traits

**DOI:** 10.1101/2025.01.13.632820

**Authors:** Xiaoqi Li, Minxing Pang, Jia Wen, Laura Y. Zhou, Laura M. Raffield, Haibo Zhou, Huaxiu Yao, Can Chen, Quan Sun, Yun Li

## Abstract

Genetic prediction of complex traits, enabled by large-scale genomic studies, has created new measures to understand individual genetic predisposition. Polygenic Risk Scores (PRS) offer a way to aggregate information across the genome, enabling personalized risk prediction for complex traits and diseases. However, conventional PRS calculation methods that rely on linear models are limited in their ability to capture complex patterns and interaction effects in high-dimensional genomic data. In this study, we seek to improve the predictive power of PRS through applying advanced deep learning techniques. We show that the Variational AutoEncoder-based model for PRS construction (VAE-PRS) outperforms currently state-of-the-art methods for biobank-level data in 14 out of 16 blood cell traits, while being computationally efficient. Through comprehensive experiments, we found that the VAE-PRS model offers the ability to capture interaction effects in high-dimensional data and shows robust performance across different pre-screened variant sets. Furthermore, VAE-PRS is easily interpretable via assessing the contribution of each individual marker to the final prediction score through the SHapley Additive exPlanations (SHAP) method, providing potential new insights in identifying trait-associated genetic variants. In summary, VAE-PRS presents a novel measure to genetic risk prediction by harnessing the power of deep learning methods, which could further facilitate the development of personalized medicine and genetic research.

## Main

The potential of genetic data in predicting complex phenotypic traits and supporting clinical decision-making has been increasingly recognized. In the past two decades, genome-wide association studies (GWAS) have substantially improved in power thanks to sample size increases in large-scale genetic studies, leading to the identification of hundreds of thousands genetic markers associated with complex traits. These findings hold the promise to estimate personalized genetic risks for complex diseases and individualized genetic prediction of complex traits. As a quantification, polygenic risk scores (PRS) summarize individuals’ risks for diseases/traits based solely on genetic profiles before disease onset. It is generally constructed with linear models based on GWAS summary statistics from a discovery population and applied to individual-level genotype data for any given target individual. PRS has shed light on how one’s genotype data contribute to a disease/trait and shows promising clinical utility^1,2^. Recent studies, such as the eMERGE study^3^ and the PRIMED consortium^4^, discussed the benefits of such integrated genetic risk assessment in routine healthcare, influencing preventive care and personalized medicine^3–6^.

With the desire to better predict phenotypic outcomes (e.g., disease risk or values from laboratory assays that are aggressive or expensive to measure) based on genetic predisposition, considerable research has been conducted to construct PRS beyond being the weighted sum of genotypes across relevant genetic variants to improve its overall predictability. Many methods have been introduced, including approaches based on pruning/clumping + thresholding (P/C+T) (e.g., PRSice and PRSice2), penalized regression (e.g. lassosum), and Bayesian methods (e.g. LDpred, PRS-CS)^7–11^. Though some of these methods have shown improved prediction accuracy, they still assume linear models without considering the complex non-linear effects among the genetic variants. Most recently, there have been various attempts to extend PRS construction methods to non-linear models^12,13^. Deep learning methods, although successfully adopted in many genomic contexts, are still under-explored for PRS construction^14,15^. Variational autoencoders (VAE), as a conventional unsupervised deep learning framework, have been successfully applied to a wide range of problems, including learning biologically meaningful latent features from cancer transcriptome and low-dimensional representation and visualization of scRNA-seq data^16,17^. It has also been used in supervised tasks such as brain age analysis^18^. In this study, we aim to improve polygenic predictions for 16 blood cell traits using VAE. Specifically, we propose VAE-PRS, a novel model which leverages a VAE-based regression framework for polygenic quantitative trait predictions (**Figure 1**) and predict PRS with supervised regression by learning the latent space of genotype data with a generative probabilistic model. Conceptually, VAE-PRS differs from existing PRS construction methods in that it simultaneously considers the prediction of the phenotypic outcome (the primary goal in PRS construction) and the reconstruction of the genotype matrix. Although auxiliary for PRS construction purposes, the latter leads to robust inference, even with correlated variants, and thus can enhance PRS construction. This design enables the model to capture complex patterns in the genetic data while maintaining a tractable computational complexity.

**Figure 1.**
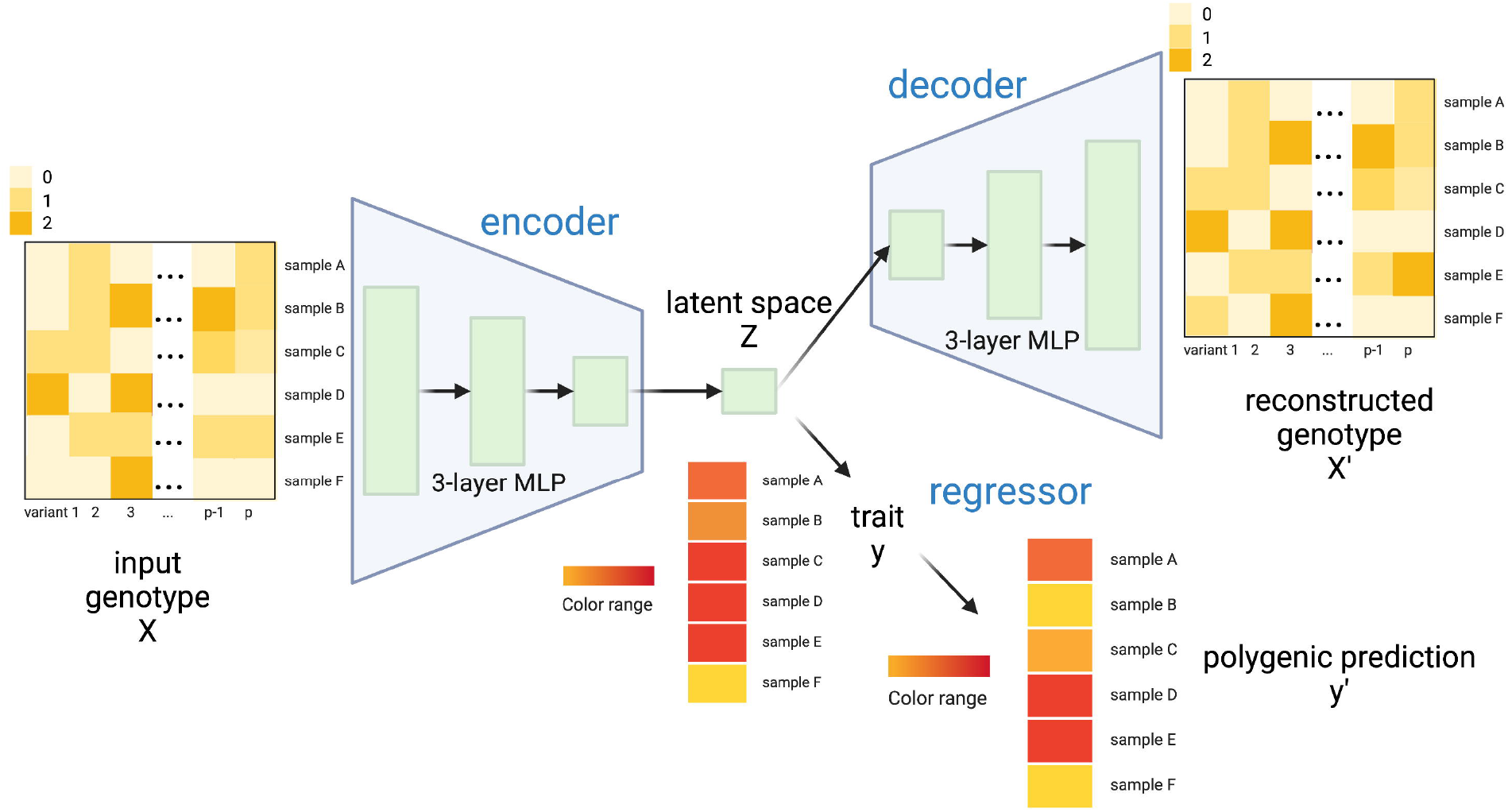
Illustration of the VAE-PRS model architecture. Our VAE-PRS architecture consists of a 3-layer MLP-based encoder and decoder, along with a separate regressor operating on the latent space. X corresponds to the input genotype: each line is a sample and each column a genetic variant; X’ is the reconstructed genotype matrix; y stands for the observed or actually measured phenotypic trait; y’ is the predicted trait based on genetic profile.

Our VAE-PRS model is composed of a 3-layer multilayer perceptron (MLP)-based encoder, a decoder, and a separate regressor operating on the latent space. The 3-layer MLP-based encoder compresses the high-dimensional input genotype data (X in **Figure 1**) into a lower-dimensional latent space, enabling the model to learn a non-linear representation of the data and complex genetic architecture underlying complex traits. The decoder, which is also a 3-layer MLP, reconstructs the input genotype matrix from the latent representation generated by the encoder. This process allows the VAE-PRS model to learn an efficient representation of the data that captures information relevant to the trait of interest. The separate regressor is designed to predict quantitative trait values assuming a Gaussian distribution learned from the latent space conditioning on the input trait (y in **Figure 1**). This approach leverages the power of the VAE framework to model the latent distribution of genotype matrix by reconstructing a new genotype matrix while explicitly modeling the relationship between the latent space and trait values (**Supplemental Methods**). This combination results in a more accurate and robust prediction of polygenic traits.

Utilizing genotype data from ∼396k European-ancestry individuals with blood cell trait information in the UK Biobank, we split them into ∼355k for training/validation (90/10 split) and ∼41k as the hold-out test set, and trained VAE-PRS models with early stopping to avoid overfitting. The quantitative blood cell traits were covariates-adjusted and inverse-normalized (**Supplemental Methods**). We compared the performance of our VAE-PRS model mainly with elastic net (EN) with the same parameters as a previous study^12^ where EN was shown to achieve the best performance among other machine-learning-based PRS methods, including MLP and CNN. We also compared C+T implemented in PRSice^7^ as a benchmark. Note that while EN and VAE-PRS typically conduct feature selection before actually running the models, PRSice employs all GWAS variants to perform clumping and thresholding with different combinations of tuning parameters in a grid search manner. For our VAE-PRS, we choose to utilize the top 100k GWAS significant variants (v100k), as using genome-wide variants is computationally infeasible. Due to the inability of EN to handle v100k variants, we further performed LD pruning and p-value thresholding at 1e-6 (abbreviated as v100k.P.T) to reduce the number of input variants (**Supplemental Methods, Table S1**). We also added VAE-PRS with v100k.P.T. variants in the comparison.

We focused on nine blood cell traits, including three white blood cell indices (eosinophil count [EOS], lymphocyte count [LYM], and white blood cell count [WBC]); three red blood cell indices (hemoglobin concentration [HGB], mean corpuscular volume [MCV], and red blood cell count [RBC]); and three platelet traits (mean platelet volume [MPV], platelet count [PLT], and plateletcrit [PCT]). Our results show that VAE-PRS using v100k variants outperforms EN and PRSice for all the nine traits, leading to 8.8-15.7% increase in Pearson correlation coefficient (PCC) between PRS and the measured phenotypes **(Figure 2, Figure S1, Table S2**). These results suggest that VAE-PRS is superior to currently the best polygenic prediction in large biobank data for blood cell traits^12^. We also observed that VAE-PRS using v100k.P.T. variants performs similarly to EN utilizing the same variant sets, but is inferior to VAE-PRS using v100k variants, likely due to the loss of information. This also suggests that the pruned correlated variants could still contribute to the trait prediction, where their effects (including potential interaction effects with other variants) could be captured by VAE-PRS, resulting in an enhanced performance.

**Figure 2.**
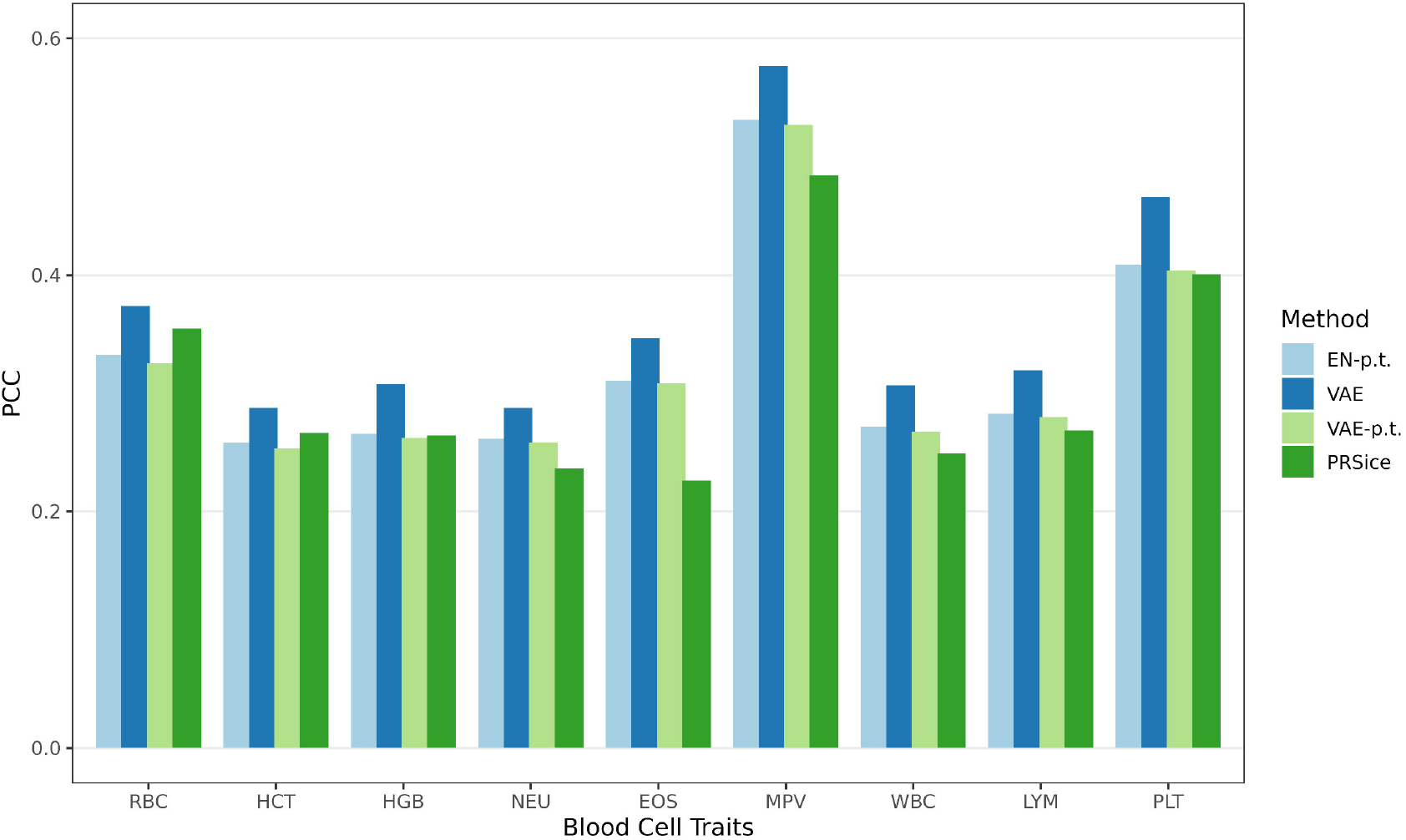
Model performance comparison for blood cell traits on ∼355k European-ancestry individuals from UK Biobank. Methods were labeled as follows: EN-p.t. (Elastic Net with v100k.P.T), VAE (VAE-PRS with v100k), VAE-p.t. (VAE-PRS with v100k.P.T) and PRSice (PRSice using all GWAS variants from the entire genome with grid search for tuning parameter optimization). The y-axis is showing the Pearson correlation between predicted values and the measure blood cell traits.

We also investigated the performance of VAE-PRS compared to EN^12^ and PRSice^7^ across different sample sizes, as many machine learning methods could only benefit with large enough samples.

Specifically, we considered sample sizes of 10k, 50k, 100k, 150k, 200k, and 355k, again, based on European ancestry individuals from UKB (**Suppelemental Methods**). All the methods were evaluated in the same held-out ∼41k individuals with the same sets of variants as employed previously. The results show that VAE-PRS exhibits its advantage over these alternative methods for larger sample sizes (>= 150k, **Figure 3**). For example, for LYM with the sample size of 150k, VAE-PRS has a PCC 17% higher than EN and 21.7% higher than PRSice. On the contrary, for sample sizes between 50k and 100k, both EN and VAE-PRS using v100k.P.T perform better than VAE-PRS with v100k for 8 and 14 out of the 16 traits, respectively. Interestingly, for a sample size of 10k, PRSice shows the best performance, where the deficiency of VAE-PRS and EN is likely due to insufficient sample size for accurate inference of the large number of parameters. This speculation is further supported by the results where pruning and thresholding largely led to improved performance for VAE-PRS and EN when sample sizes are below 150k and above 10k (**Figures 3, Table S3, Figure S1**). These results emphasize the importance of feature selection or dimension reduction for deep learning methods when sample sizes are below 100,000 and highlight the potential of the VAE-PRS model for polygenic trait prediction in large-scale biobank-level datasets. On the other hand, it also confirms that sample size is still a bottleneck for current deep learning methods to advance in genetic studies.

**Figure 3.**
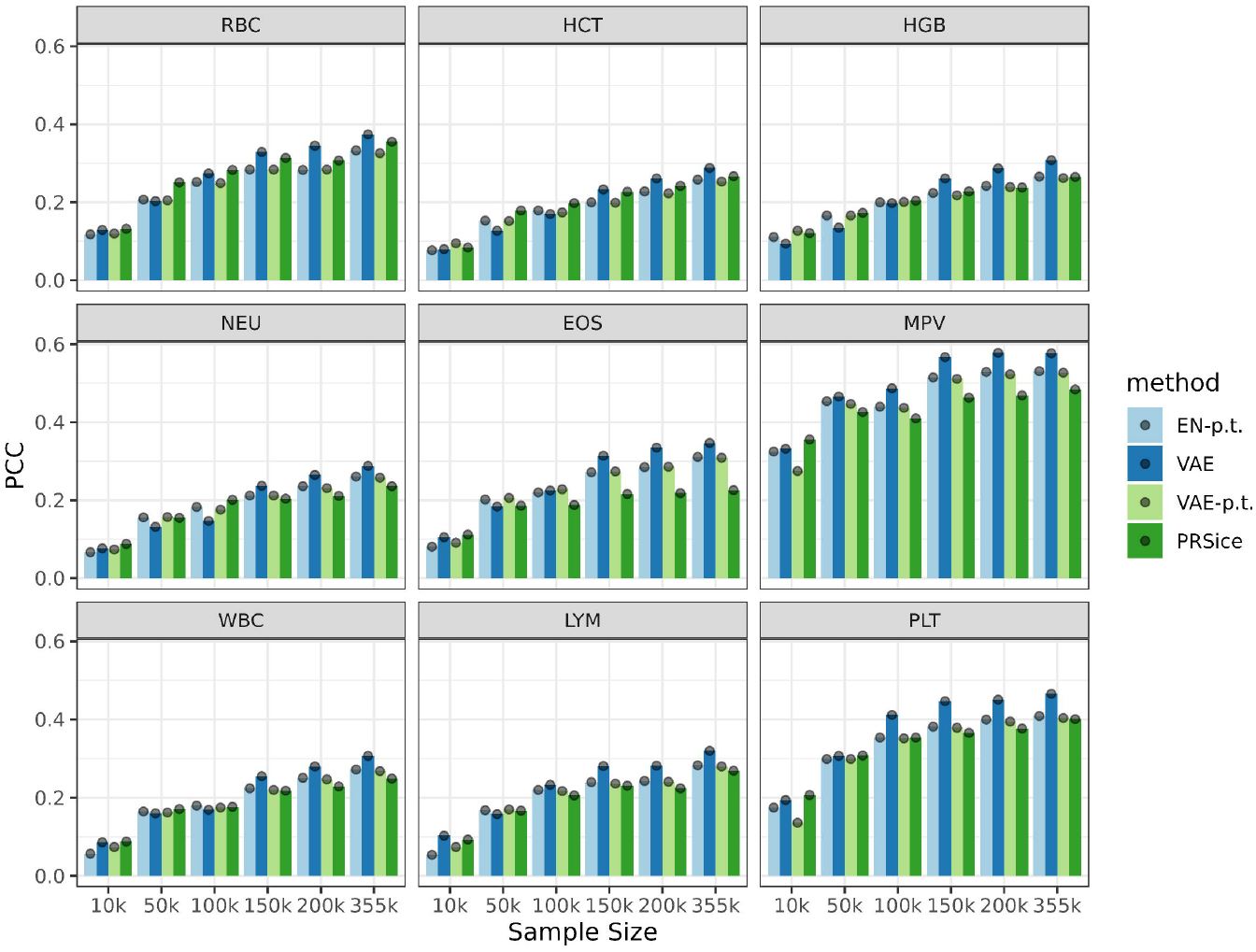
Minimum sample size needed to benefit from VAE. We conducted assessments across varying sample sizes: 10k, 50k, 100k, 150k, 200k, and 355k. Methods were labeled as follows: EN-p.t. (Elastic Net using v100k.P.T), VAE (VAE-PRS using v100k), VAE-p.t. (VAE-PRS using v100k.P.T) and PRSice (PRSice using all GWAS variants from the entire genome with grid search for tuning parameter optimization).

As mentioned previously, VAE-PRS utilizing v100k.P.T. variants performs less well than VAE-PRS with v100k variants, which is again confirmed in **Figure 3**, especially with larger sample size (>= 150k). For example, with n = 200K, VAE-PRS with v100k (VAE in **Figure 3**) achieves a PCC of 0.345 for RBC, while for VAE-PRS with v100k.P.T. (VAE.p.t in **Figure 3**), PCC decreases to 0.284. This observation suggests that pruning and thresholding may lead to information loss. To further assess the performance of our VAE-PRS model on different variant sets, we employed carefully selected sets of variants generated from conditional analyses (CA) by a previous study where such strategy was shown to have a better predictive performance for blood cell traits^12,19^. We found that both EN and VAE-PRS demonstrated improved performance with the carefully selected variants. On average, EN shows 20.1% improvement in terms of PCC, while VAE-PRS shows 10.2% (**Figure S2, Table S4**). With fewer number of input variants, the advantages of VAE-PRS are less pronounced. However, selecting CA variants demands significant time and efforts^20^. VAE-PRS would be valuable when no carefully selected variants are available for PRS analysis.

To better understand the contribution of individual markers, we computed the SHapley Additive exPlanations (SHAP) feature importance scores^21^. These scores quantify the contribution of each variant to the overall model by dropping each one at a time and calculating the prediction difference. We compared SHAP scores with GWAS p-values and the absolute value of the GWAS effect size estimates for MPV in **Figure 4** as an example, and for all the other traits in **Figure S3 a-o**. For all the blood cell traits where VAE-PRS exhibit high predictive performances, we observed that the overall correspondence between SHAP scores and GWAS p-values (or effect sizes) are relatively low, though some high GWAS significance loci could be captured by SHAP values. For example, PCC and Spearman correlation for MPV (**Figure 4**) is 0.22 and 0.09 between SHAP and GWAS p-values (−0.07 and −0.30 between SHAP and GWAS effect sizes), similarly for other traits (**Table S5**). The relatively low agreement suggests that variants contribute to VAE-PRS predictions in an order not necessarily determined by GWAS significance or magnitude of effect, which may potentially aid us in re-prioritization and/or combination of variants. For example, variant 3:143021856:G:C (hg19, rs9842051) on chromosome 3 (highlighted with the red box on the left in **Figure 4**) is ranked 77th in terms of SHAP feature importance scores, but is only ranked 78,088th for GWAS p-values and 44,143th for GWAS effect sizes.

**Figure 4.**
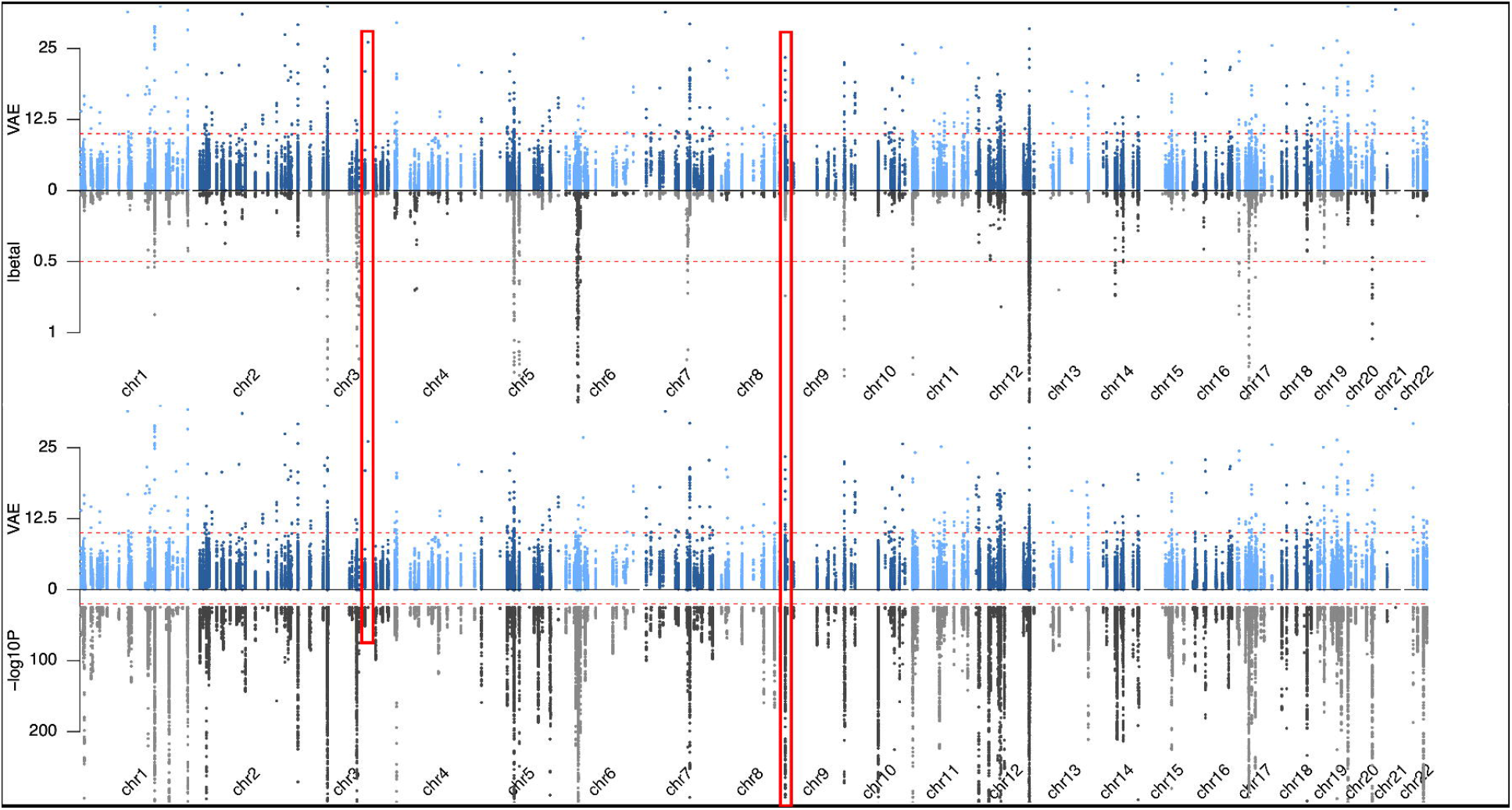
Feature importance mirror plots for MPV. The upper mirror plot compares SHAP values with absolute values for GWAS effect sizes (|beta|), while the lower mirror plot compares SHAP with GWAS p-values (−log10).

We then proceeded with explorations for variants with the highest SHAP scores, assessing potential the interaction effects between variants. Specifically, we took MPV as an example and focused on the 100 variants with the top SHAP scores (top 0.1%). We evaluated the significance of pairwise interactions in the test samples using v100k variants. We discovered six marginally significant variant pairs with false discovery rates (FDR) < 0.05. The most significant pair was 9:273178:G:GA (hg19, rs3215853) and 3:143021856:G:C (hg19, rs9842051), located in the coding regions of the *DOCK8* and *SLC9A9* genes respectively (**Figure 5**; also highlighted with red boxes in **Figure 4**). We note that both of the genes have been previously reported to be significantly associated with MPV or platelet count^19,22^. Interestingly, *DOCK8* and *SLC9A9* have also been reported to be significant for tumor mutational burden infiltration (TMB-IF) models^23^. Given that high platelet counts are associated with an increased risk of cancer, this finding suggests a potential link between these genes and cancer risk. To the best of our knowledge, there has been no literature reporting the interaction between these two variants or genes before, in part because they are located on two different chromosomes. This statistically significant interaction between 9:273178:G:GA and 3:143021856:G:C not only provides insights into the genetic architecture underlying MPV, but also highlights the associations between MPV, cancer risks, and the immune system mediated by these variants or genes. Further investigations of these interactions may help improve our understanding of the complex relationships between blood cell traits, immune response, cancer development, and other mechanisms related to health and disease outcomes, ultimately leading to more targeted therapeutic strategies and improved health care.

**Figure 5.**
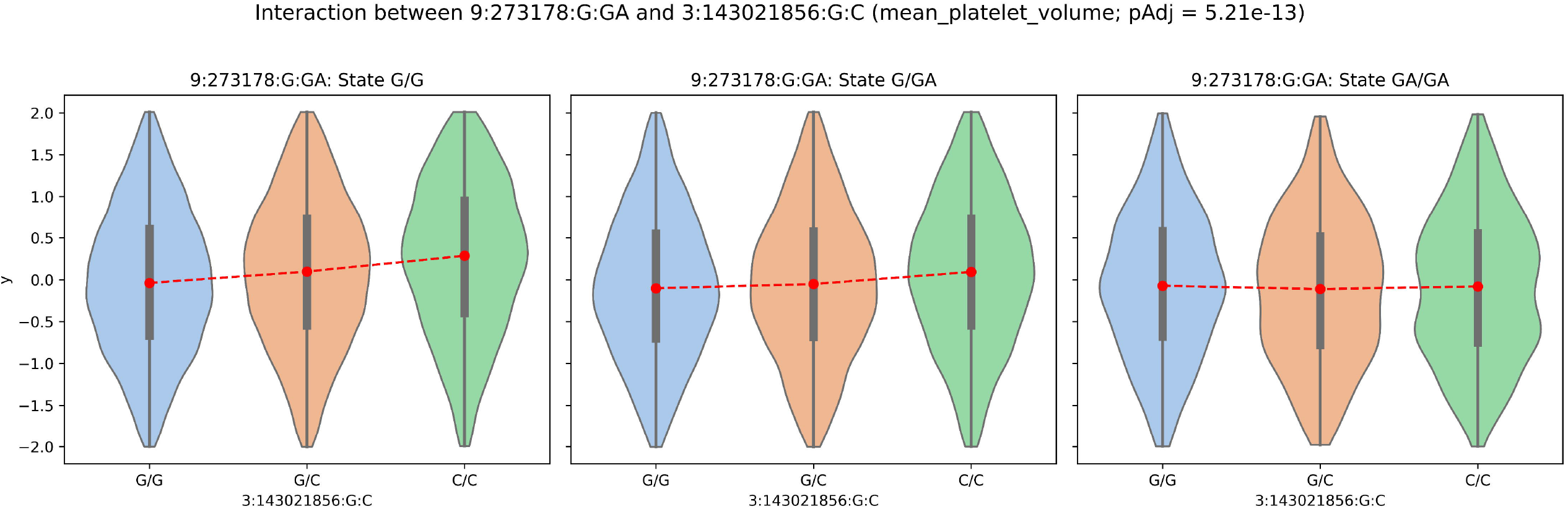
Statistically significant interaction between 3:143021856:G:C and 9:273178:G:GA for MPV (FDR = 5.21e-13). The y-axis represents measured MPV after inverse normal transformation and covariate adjustments, and the x-axis shows the genotypes of 3:143021856:G:C.The three sub-plots represent individuals whose genotypes of 9:273178:G:GA are G/G, G/GA and GA/GA from left to right. The red dashed line connects the median of the transformed and adjusted MPV values.

We also evaluated the computational efficiency of the VAE-PRS model in comparison to EN. Faced with the problems of loading the entire dataset into memory, we decided to take advantage of batch training and employed early stopping to avoid model overfitting. VAE-PRS using v100k.P.T. demands only about half of the computational time and ∼1/30 of the memory usage compared to EN trained with the same variants (**Figure S4, Tables S6-7**). While VAE-PRS with v100k variants requires more computation time, it exhibits a substantial improvement in performance (12.1% on average for PCC) and consumes only ∼1/15 memory compared to EN using only v100k.P.T. variants. VAE-PRS maintains an advantage in terms of memory usage even when using 19-172 times more variants, indicating its utility to be applied effectively for large-scale genetic studies.

While VAE-PRS demonstrates promising results for polygenic trait prediction, there are some aspects that could be further improved in future studies. First, due to current computational limitations, we only utilized 100k variants as inputs for model training, although the number of variants included was much larger compared to EN. Future efforts are warranted to perform model training with more variants in consideration after computational breakthrough. Second, like many other PRS methods, we did not provide any uncertainty measurements of individual PRS estimates. One could explore Bayesian VAE model^24^ for this task, which requires comprehensive evaluations in future studies. Third, we primarily focused on individuals with European ancestry in this study, as other ancestry groups have insufficient sample sizes to benefit from VAE-PRS models. Although transfer learning techniques could be adopted with models trained on large-scale European individuals, our preliminary results suggest that only 9/15 traits showed slightly better performance via transfer learning compared to directly applying European-trained models (**Table S8**). We hypothesize this is in part due to the tuning sample size limitation, which is also observed in our sample size evaluations (**Figure 3, Figure S1)**. These findings highlight the persistent challenges of developing broadly generalizable PRS methods across diverse populations with deep learning models. Last, our VAE-PRS framework needs access of individual-level data for model training, which may be limited by availability. However, with efforts allowing for broad data access and sharing, such as NIH’s All of Us program, we anticipate more individual data will be available for PRS model training to benefit from deep learning models.

In summary, we here present VAE-PRS, which leverages the power of VAE and allows us to capture complex interactions between genetic markers and to improve the accuracy of polygenic trait prediction for blood cell traits. By training in parallel, VAE-PRS is computationally more efficient compared to alternative methods with no or negligible sacrifice of prediction accuracy. Our results have unveiled the potential of harnessing the power of deep learning models, specifically VAE, to facilitate the development of polygenic risk prediction. We also provided recommendations on the sample sizes from empirical results about when to use VAE-PRS models. In the future, with increased sample sizes of large-scale genomic studies, we anticipate greater benefits of VAE-PRS.

## Acknowledgement

This study was supported by NIH grant U01HG011720 and R01HL146500. In addition, QS was also supported by U24AR076730, JW by T32ES007018, YL by R01HL165061. This research has been conducted using the UK Biobank Resource under Application Number 25953.

## Author Contributions

Study design and conceptualization: XL, MP, QS, YL; analysis: XL, MP, QS, JW, LYZ; data generating and coordinating: LMR, YL; manuscript writing: XL, QS, YL; supervision: QS, YL. All authors contributed to manuscript revisions.

## Data and Code Availability

The scripts used in this article have been made public on: https://github.com/ahalxq/vaeprs. Our study used data from UK Biobank, which we applied for and downloaded from https://www.ukbiobank.ac.uk. The UK Biobank has ethics approval from the North West Multi-centre Research Ethics Committee (MREC).

## Web resources

UK Biobank: https://www.ukbiobank.ac.uk

PLINK Pipelines: https://github.com/arnor-sigurdsson/plink_pipelines

EN: https://github.com/xuyu-cam/PGS-BC-Traits-Using-ML-DL

PRSice2: https://choishingwan.github.io/PRSice/

PLINK2: https://www.cog-genomics.org/plink/2.0/

## References

1 Lewis, C.M., and Vassos, E. (2020). Polygenic risk scores: from research tools to clinical instruments. Genome Medicine 12, 44. 10.1186/s13073-020-00742-5.

2 Kumuthini, J., Zick, B., Balasopoulou, A., Chalikiopoulou, C., Dandara, C., El-Kamah, G., Findley, L., Katsila, T., Li, R., Maceda, E.B., et al. (2022). The clinical utility of polygenic risk scores in genomic medicine practices: a systematic review. Hum Genet 141, 1697–1704. 10.1007/s00439-022-02452-x.

3 Linder, J.E., Allworth, A., Bland, S.T., Caraballo, P.J., Chisholm, R.L., Clayton, E.W., Crosslin, D.R., Dikilitas, O., DiVietro, A., Esplin, E.D., et al. (2023). Returning integrated genomic risk and clinical recommendations: The eMERGE study. Genet Med 25, 100006. 10.1016/j.gim.2023.100006.

4 Kachuri, L., Chatterjee, N., Hirbo, J., Schaid, D.J., Martin, I., Kullo, I.J., Kenny, E.E., Pasaniuc, B., Polygenic Risk Methods in Diverse Populations (PRIMED) Consortium Methods Working Group, Witte, J.S., et al. (2024). Principles and methods for transferring polygenic risk scores across global populations. Nat Rev Genet 25, 8–25. 10.1038/s41576-023-00637-2.

5 Pacyna, J.E., Ennis, J.S., Kullo, I.J., and Sharp, R.R. (2023). Examining the Impact of Polygenic Risk Information in Primary Care. J Prim Care Community Health 14, 21501319231151766. 10.1177/21501319231151766.

6 Pacheco, J.A., Rasmussen, L.V., Wiley, K., Person, T.N., Cronkite, D.J., Sohn, S., Murphy, S., Gundelach, J.H., Gainer, V., Castro, V.M., et al. (2023). Evaluation of the portability of computable phenotypes with natural language processing in the eMERGE network. Sci Rep 13, 1971. 10.1038/s41598-023-27481-y.

7 Choi, S.W., and O’Reilly, P.F. (2019). PRSice-2: Polygenic Risk Score software for biobank-scale data. GigaScience 8, giz082. 10.1093/gigascience/giz082.

8 Choi, S.W., Mak, T.S.-H., and O’Reilly, P.F. (2020). Tutorial: a guide to performing polygenic risk score analyses. Nature Protocols 15, 2759–2772. 10.1038/s41596-020-0353-1.

9 Ge, T., Chen, C.-Y., Ni, Y., Feng, Y.-C.A., and Smoller, J.W. (2019). Polygenic prediction via Bayesian regression and continuous shrinkage priors. Nature Communications 10, 1776. 10.1038/s41467-019-09718-5.

10 Mak, T.S.H., Porsch, R.M., Choi, S.W., Zhou, X., and Sham, P.C. (2017). Polygenic scores via penalized regression on summary statistics. Genetic Epidemiology 41, 469–480. 10.1002/gepi.22050.

11 Privé, F., Arbel, J., and Vilhjálmsson, B.J. (2020). LDpred2: better, faster, stronger. Bioinformatics 36, 5424–5431. 10.1093/bioinformatics/btaa1029.

12 Xu, Y., Vuckovic, D., Ritchie, S.C., Akbari, P., Jiang, T., Grealey, J., Butterworth, A.S., Ouwehand, W.H., Roberts, D.J., Di Angelantonio, E., et al. (2022). Machine learning optimized polygenic scores for blood cell traits identify sex-specific trajectories and genetic correlations with disease. Cell Genomics 2, 100086. 10.1016/j.xgen.2021.100086.

13 Elgart, M., Lyons, G., Romero-Brufau, S., Kurniansyah, N., Brody, J.A., Guo, X., Lin, H.J., Raffield, L., Gao, Y., Chen, H., et al. (2022). Non-linear machine learning models incorporating SNPs and PRS improve polygenic prediction in diverse human populations. Communications Biology 5, 856. 10.1038/s42003-022-03812-z.

14 Sigurdsson, A.I., Louloudis, I., Banasik, K., Westergaard, D., Winther, O., Lund, O., Ostrowski, S.R., Erikstrup, C., Pedersen, O.B.V., Nyegaard, M., et al. (2023). Deep integrative models for large-scale human genomics. Nucleic Acids Research, gkad373. 10.1093/nar/gkad373.

15 Arnór I. Sigurdsson, Kirstine Ravn, Ole Winther, Ole Lund, Søren Brunak, Bjarni J. Vilhjálmsson, and Simon Rasmussen (2022). Improved prediction of blood biomarkers using deep learning. medRxiv, 2022.10.27.22281549. 10.1101/2022.10.27.22281549.

16 Way, G.P., and Greene, C.S. (2017). Extracting a biologically relevant latent space from cancer transcriptomes with variational autoencoders. In Biocomputing 2018 (WORLD SCIENTIFIC), pp. 80–91. 10.1142/9789813235533_0008.

17 Wang, D., and Gu, J. (2018). VASC: Dimension Reduction and Visualization of Single-cell RNA-seq Data by Deep Variational Autoencoder. Genomics, Proteomics & Bioinformatics 16, 320–331. 10.1016/j.gpb.2018.08.003.

18 Zhao, Q., Adeli, E., Honnorat, N., Leng, T., and Pohl, K. (2019). Variational AutoEncoder For Regression: Application to Brain Aging Analysis. In Medical Image Computing and Computer Assisted Intervention (Springer International Publishing), pp. 823–831. 10.1007/978-3-030-32245-8_91.

19 Vuckovic, D., Bao, E.L., Akbari, P., Lareau, C.A., Mousas, A., Jiang, T., Chen, M.-H., Raffield, L.M., Tardaguila, M., Huffman, J.E., et al. (2020). The Polygenic and Monogenic Basis of Blood Traits and Diseases. Cell 182, 1214-1231.e11. 10.1016/j.cell.2020.08.008.

20 Raffield, L.M., Iyengar, A.K., Wang, B., Gaynor, S.M., Spracklen, C.N., Zhong, X., Kowalski, M.H., Salimi, S., Polfus, L.M., Benjamin, E.J., et al. (2020). Allelic Heterogeneity at the CRP Locus Identified by Whole-Genome Sequencing in Multi-ancestry Cohorts. Am J Hum Genet 106, 112–120. 10.1016/j.ajhg.2019.12.002.

21 Lundberg, S.M., and Lee, S.-I. (2017). A Unified Approach to Interpreting Model Predictions. In Advances in Neural Information Processing Systems, I. Guyon, U. V. Luxburg, S. Bengio, H. Wallach, R. Fergus, S. Vishwanathan, and R. Garnett, eds. (Curran Associates, Inc.).

22 Barton, A.R., Sherman, M.A., Mukamel, R.E., and Loh, P.-R. (2021). Whole-exome imputation within UK Biobank powers rare coding variant association and fine-mapping analyses. Nat Genet 53, 1260–1269. 10.1038/s41588-021-00892-1.

23 Xie, C., Wu, H., Pan, T., Zheng, X., Yang, X., Zhang, G., Lian, Y., Lin, J., and Peng, L. (2021). A novel panel based on immune infiltration and tumor mutational burden for prognostic prediction in hepatocellular carcinoma. Aging (Albany NY) 13, 8563–8587. 10.18632/aging.202670.

24 Daxberger, E., and Herna ndez-Lobato, J. M. Bayesian Variational Autoencoders for Unsupervised Out-of-Distribution Detection. In 4th workshop on Bayesian Deep Learning, NeurIPS 2019 10.48550/arXiv.1912.05651.

