## Supplemental Figures and Methods for "Variational Autoencoder-based Model Improves Polygenic Prediction in Blood Cell Traits"

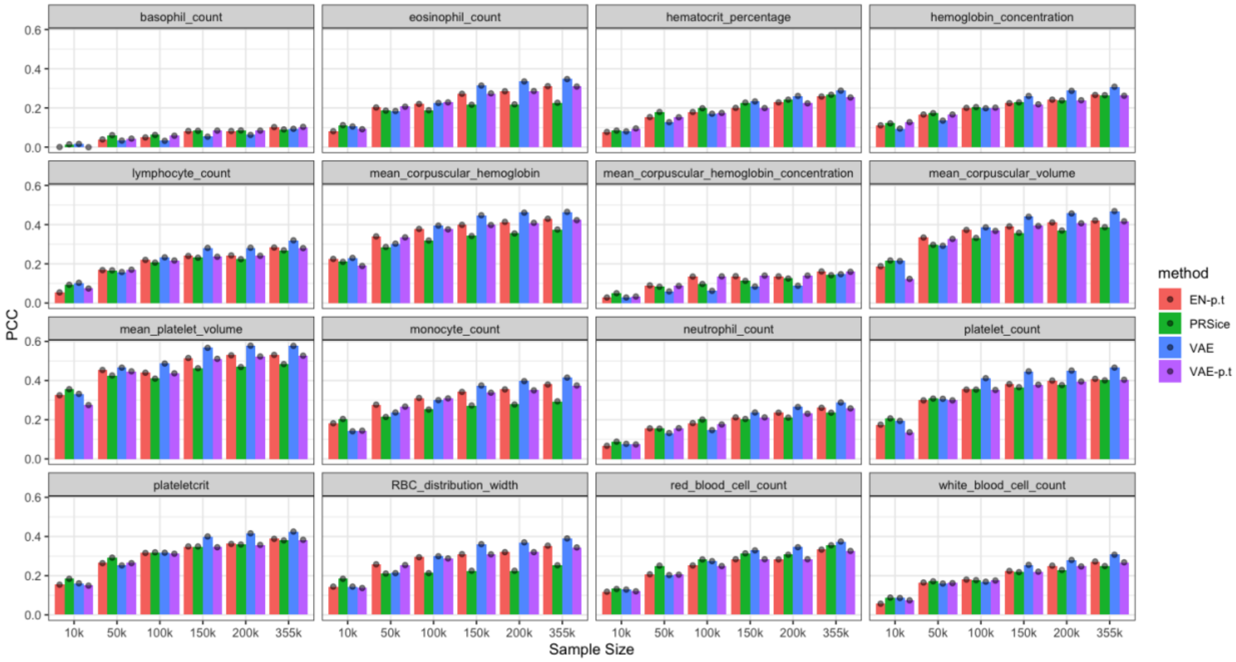


**Figure S1. Perason correlation coefficients of all the 16 blood cell trait PRS with different training sample sizes.** The labels represent: EN-p.t. (Elastic Net with v100k.P.T variants), PRSice (PRSice using all GWAS variants regardless of LD and p-value), VAE (VAE-PRS with v100k variants), and VAE-p.t. (VAE-PRS with v100k.P.T variants).

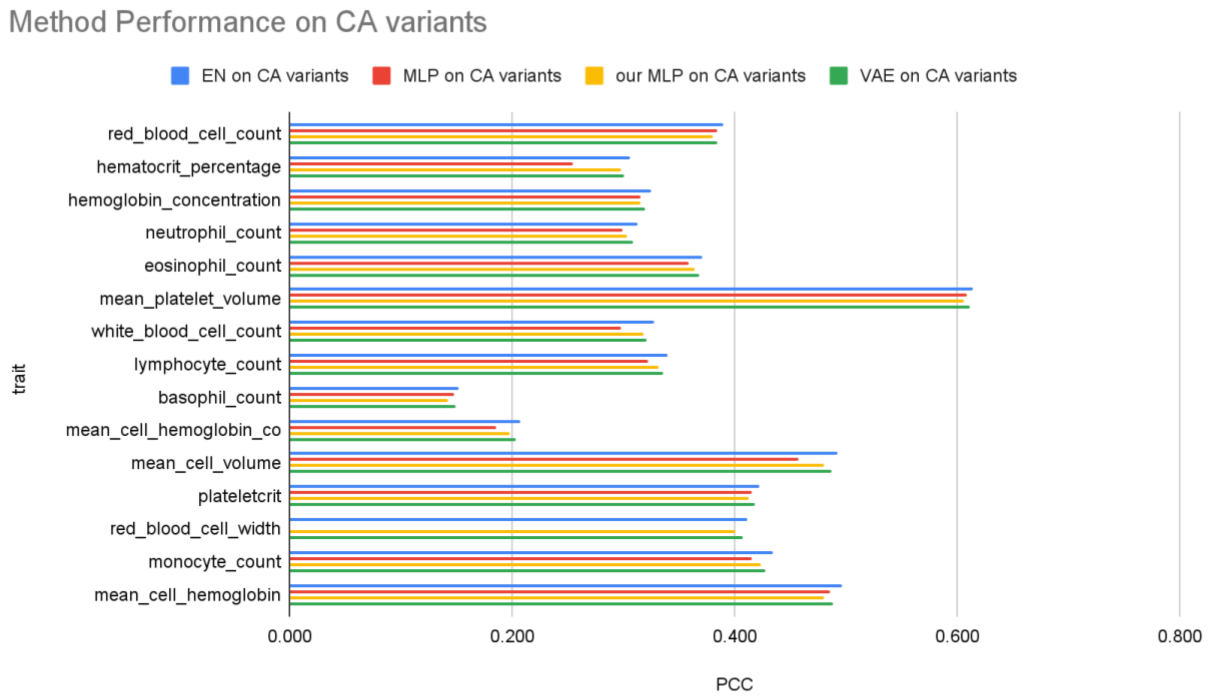


**Figure S2. Pearson Correlation Coefficients of blood cell trait PRS from different methods using carefully selected variants from conditional analysis (CA).** The x-axis denotes the Pearson Correlation Coefficient (PCC), while the y-axis represents different traits. VAE: our VAE model; EN: elastic net model; MLP: Pearson correlation coefficient from multilayer perceptron models reported in Xu et al. [9] for blood cell traits; our MLP: results from our self-implemented fine-tuned MLP models.

**(a) Eosinophil count (EOS)**

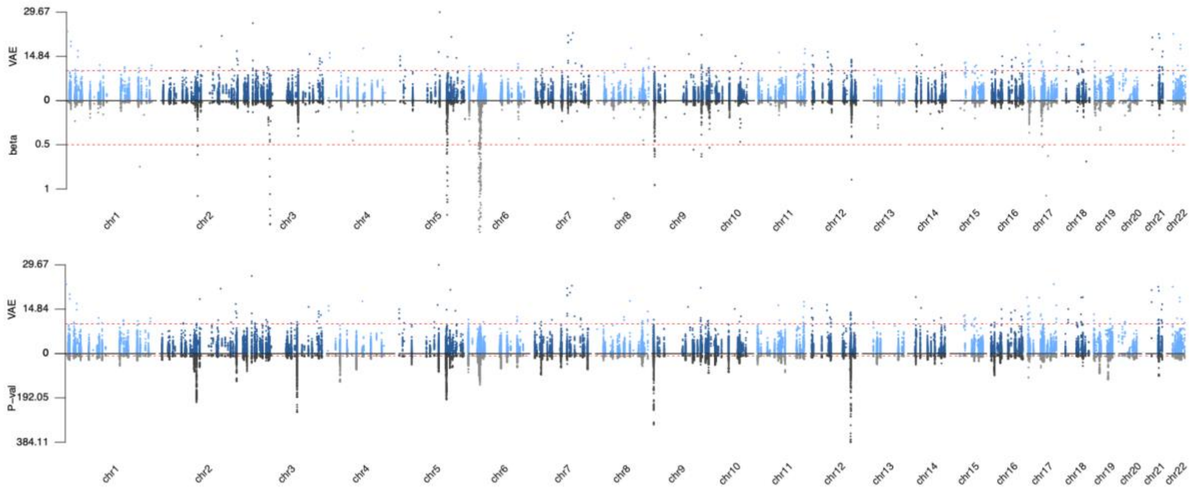


**(b) Hematocrit percentage (HCT)**

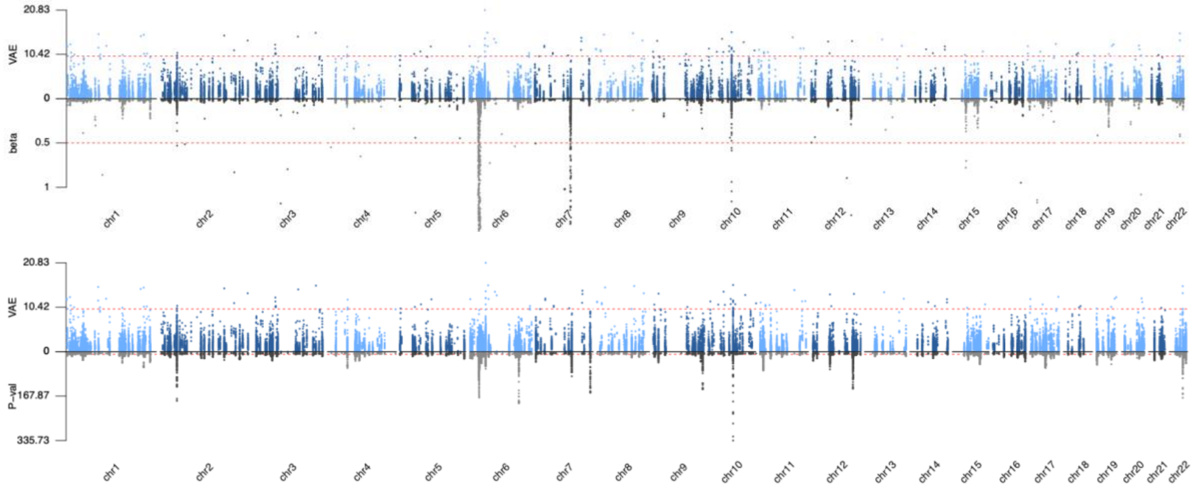


**(c) Hemoglobin concentration (HGB)**

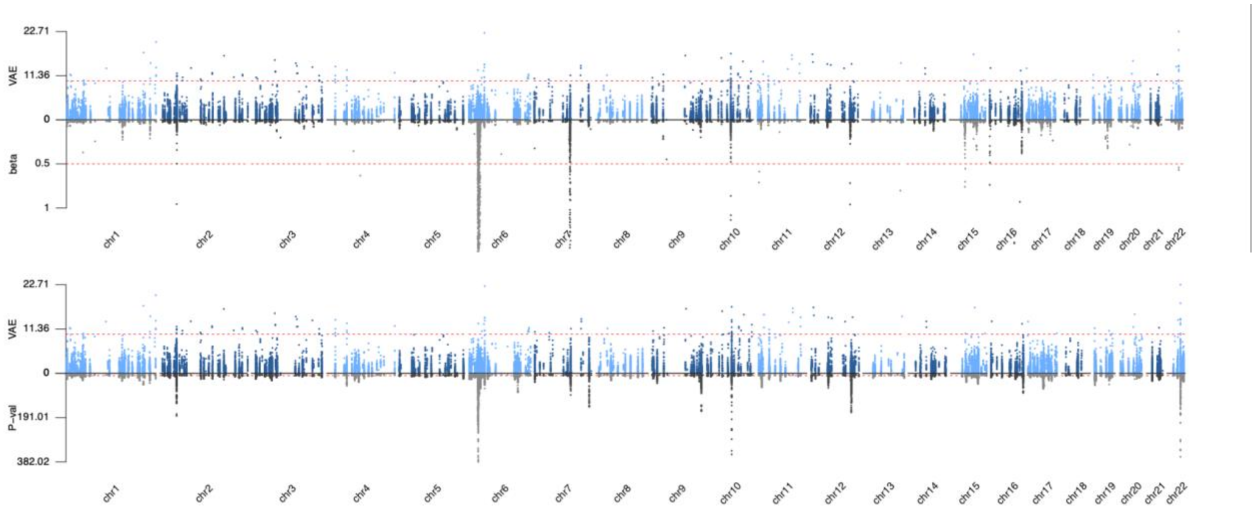

**(d) Lymphocyte count (LYM)**

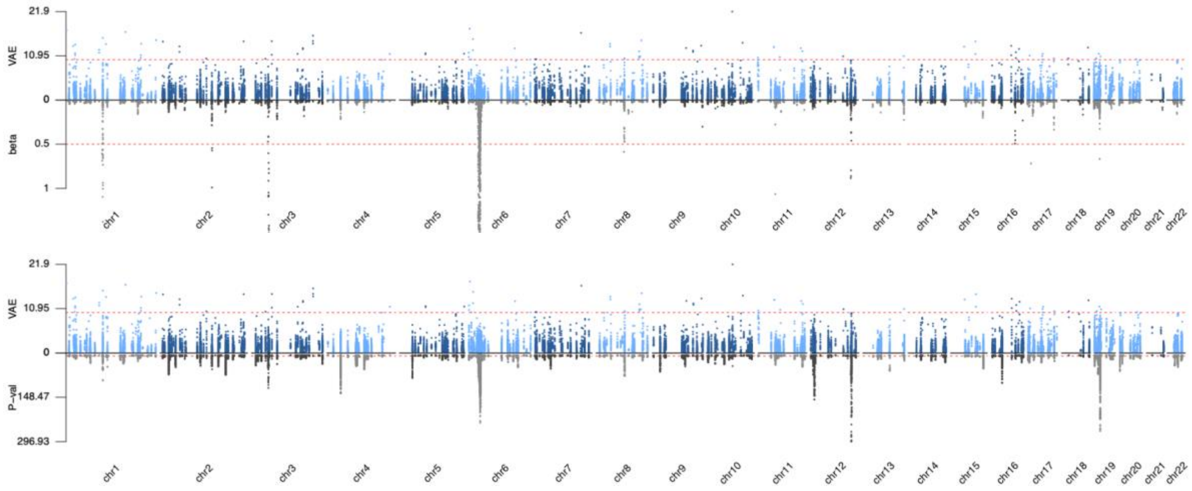


**(e) Basophil count (BASO)**

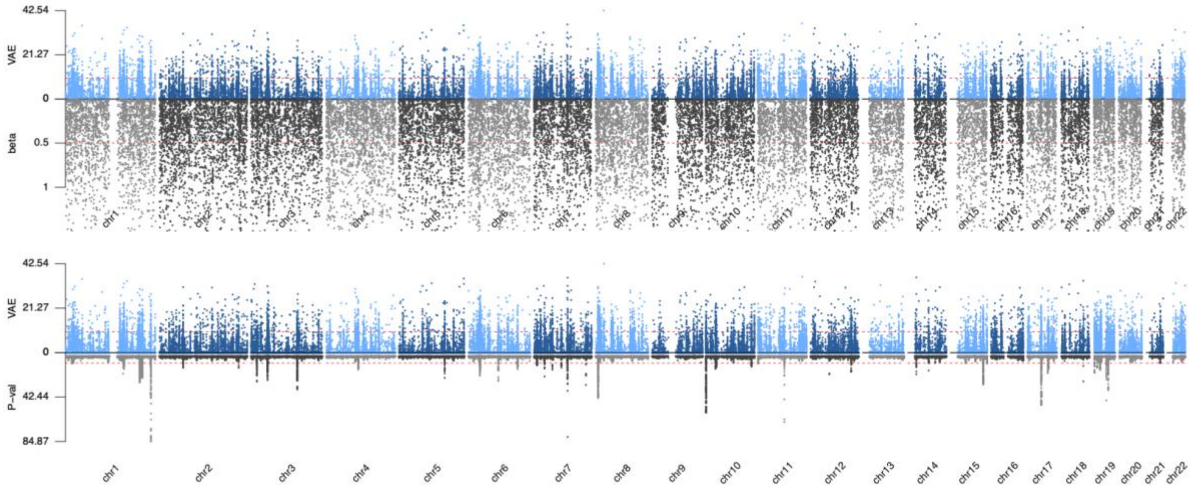


**(f) Mean corpuscular hemoglobin concentration (MCHC)**

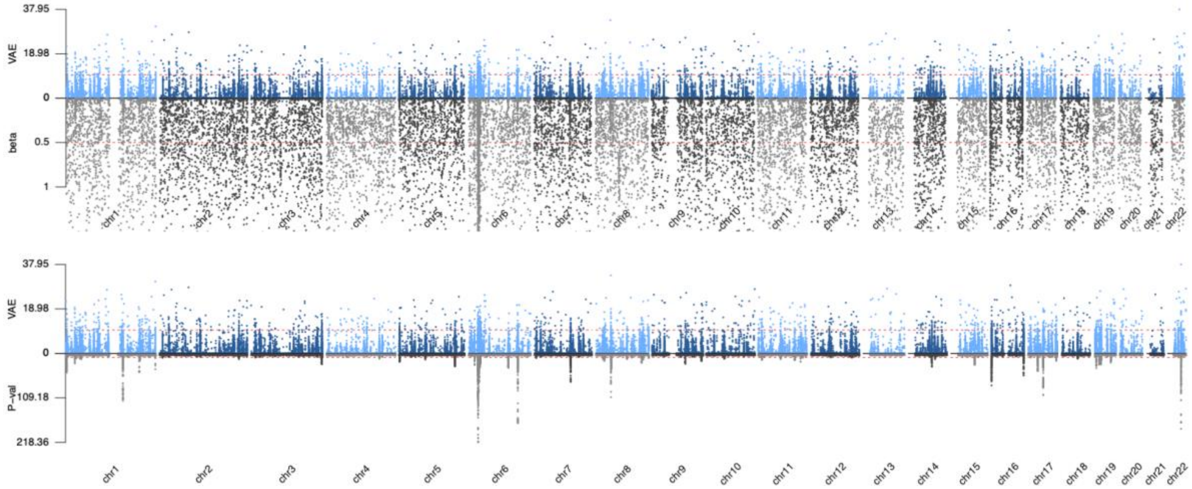


**(g) Mean corpuscular hemoglobin (MCH)**

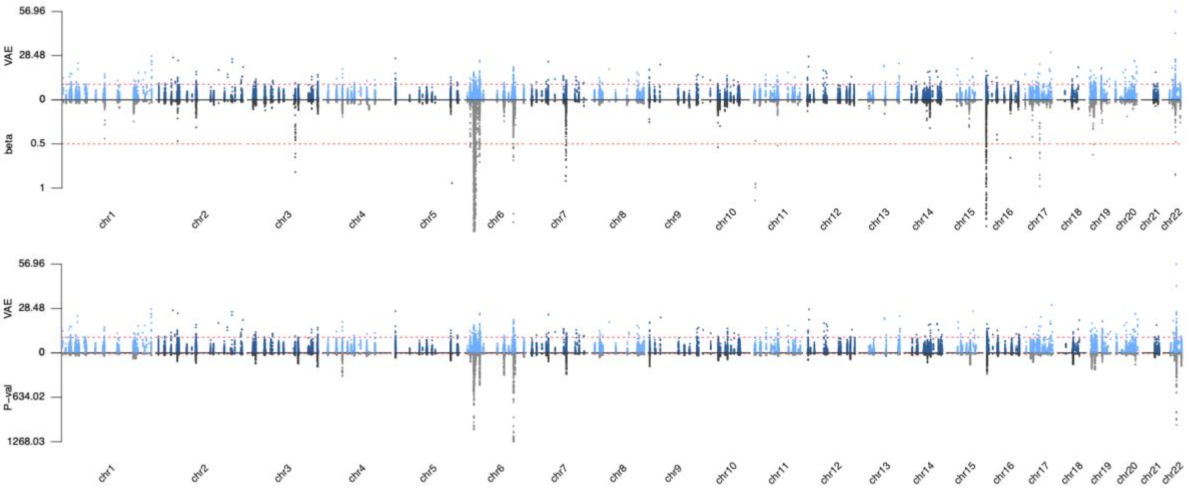


**(h) Mean corpuscular volume (MCV)**

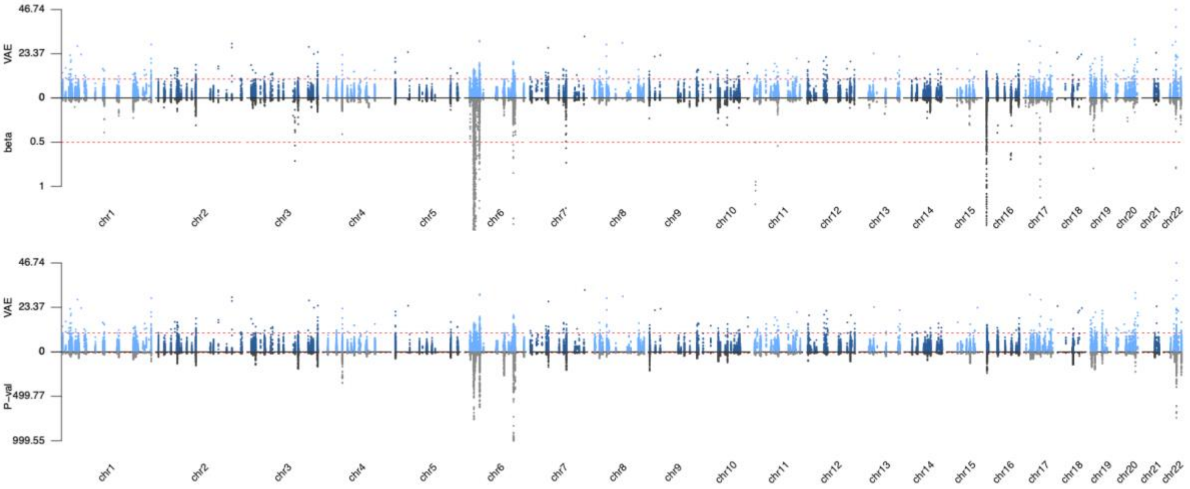


**(i) Mean platelet volume (MPV)**

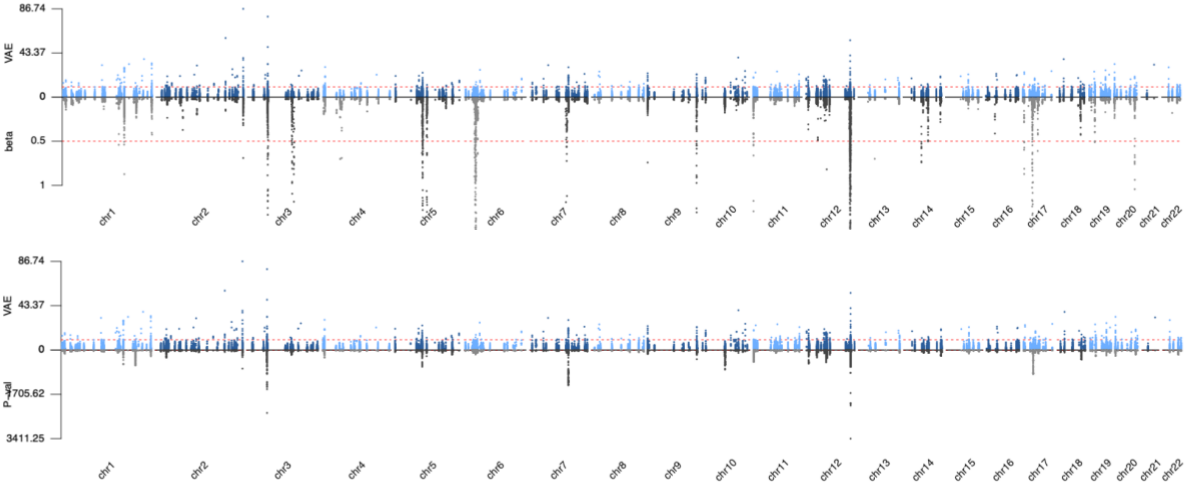


**(j) Monocyte count (MONO)**

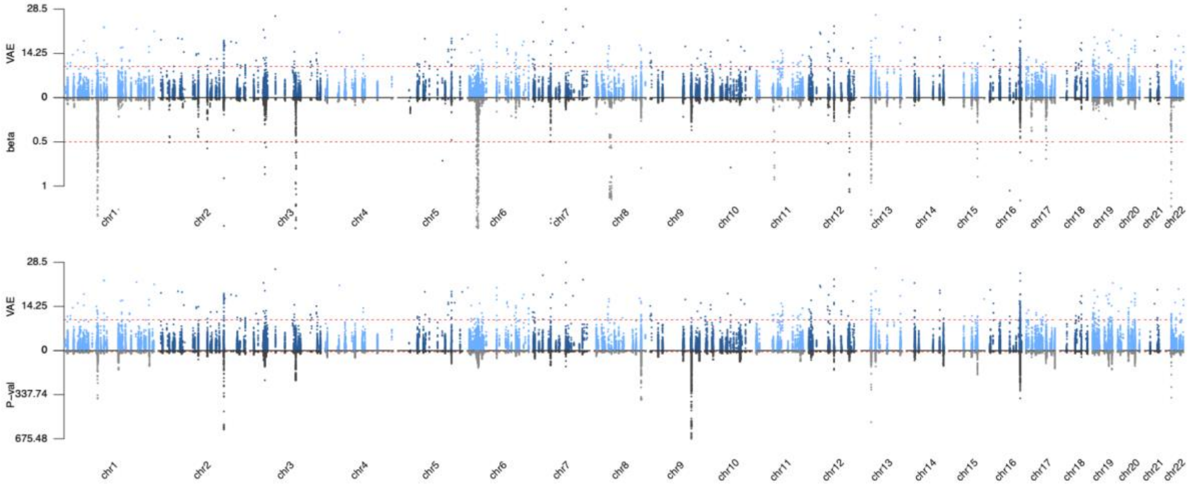


**(k) Neutrophil count (NEU)**

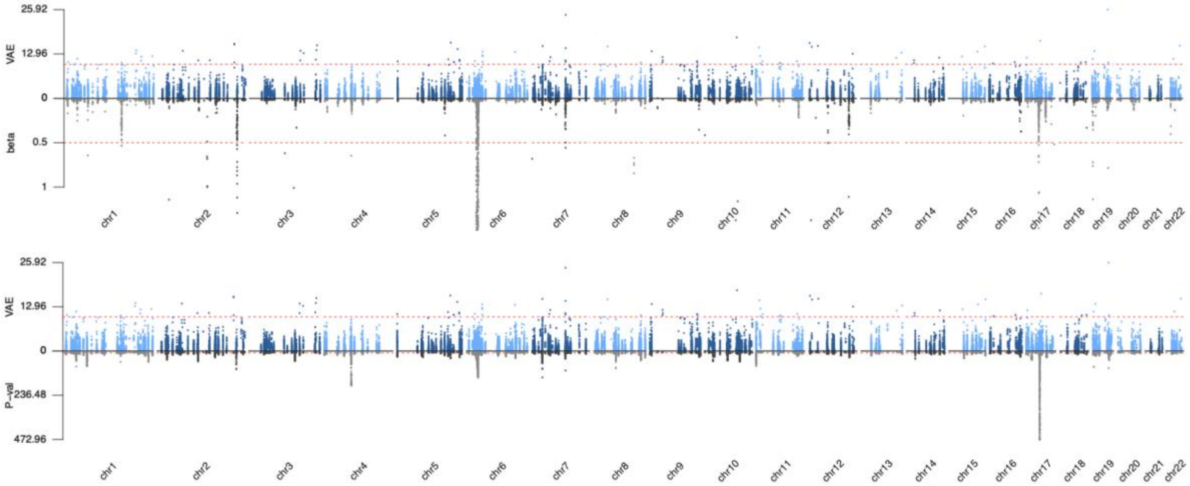


**(l) Platelet Count (PLT)**

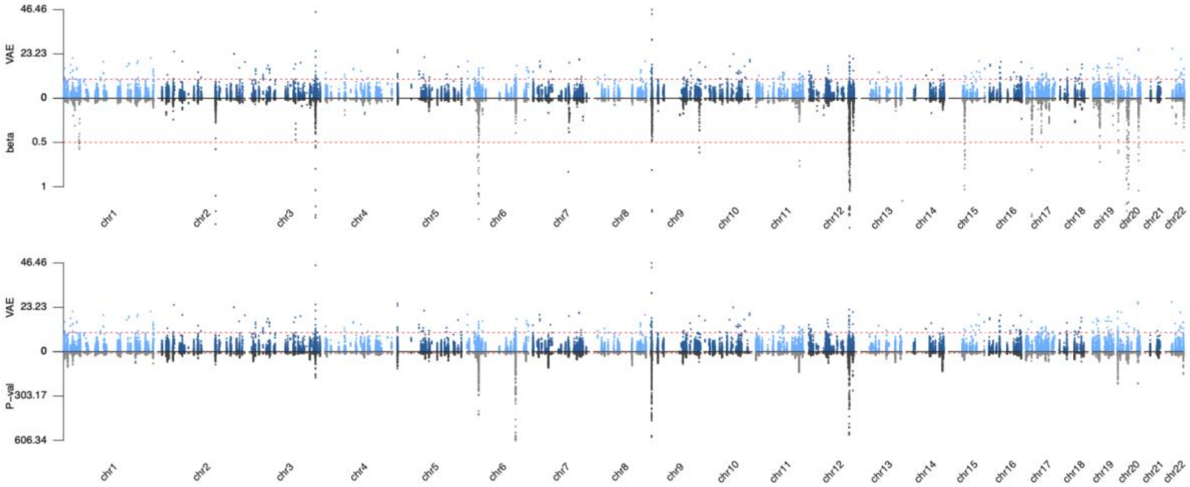


**(m) Red blood cell count (RBC)**

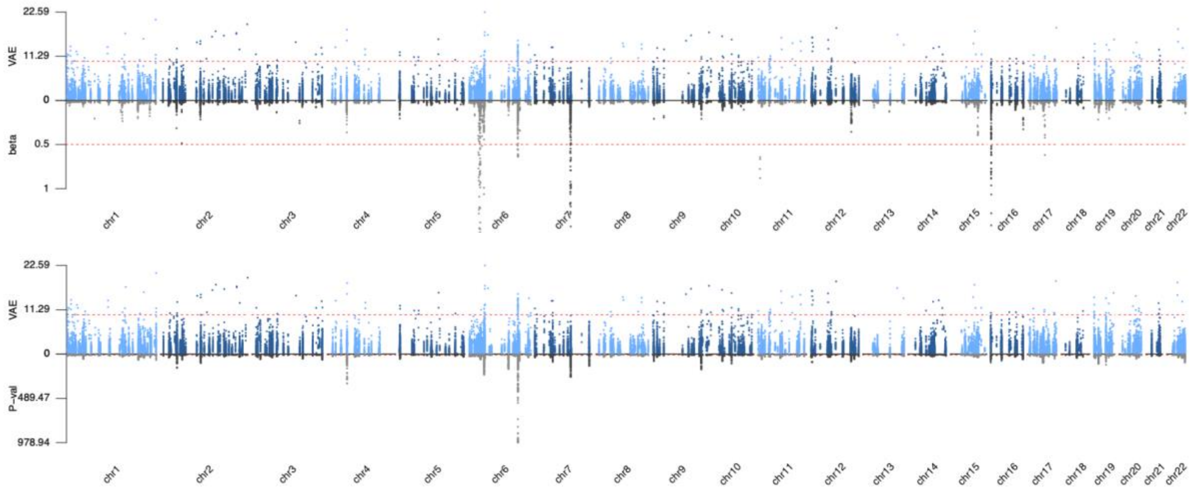


**(n) Red blood cell width (RDW)**

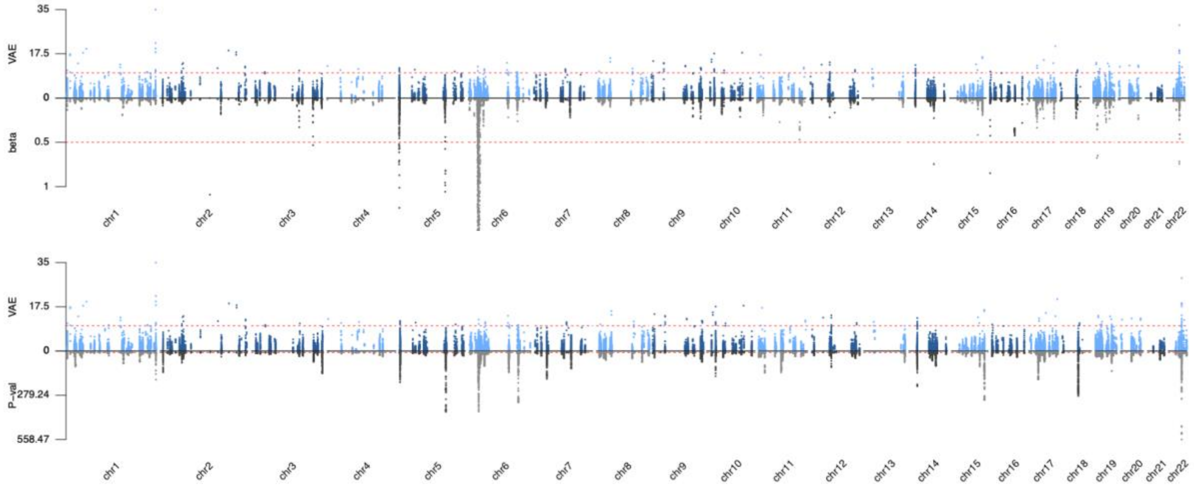


**(o) White blood cell count (WBC)**

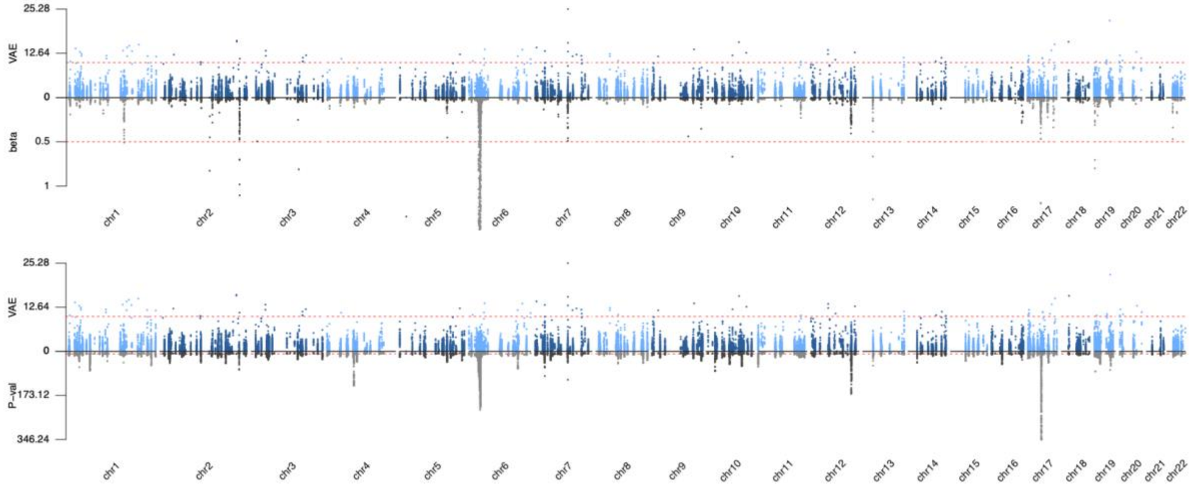


**Figure S3. Comparison between SHAP scores and GWAS effect sizes (upper) and -$\log_{10}$ transformed p-values (lower) for 15 blood cell traits.** Each figure consists of two panels representing results for each blood cell trait. The upper figure presents a comparison between the feature importance ascribed by VAE and the beta values from GWAS analysis for the top 100k variant. The lower figure contrasts the VAE feature importance with the p-values derived from GWAS analysis for the top 100k variant.

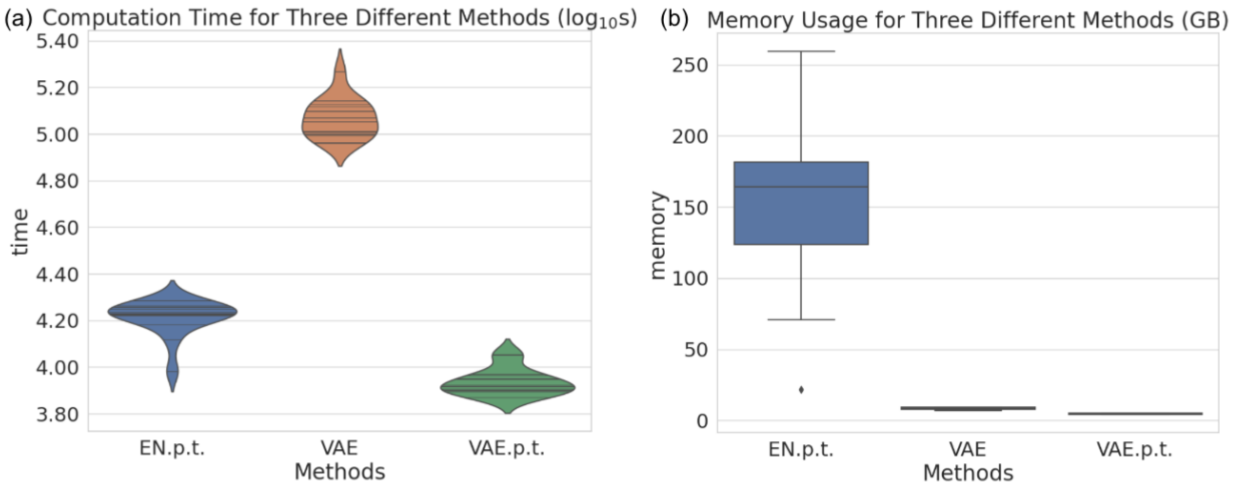


**Figure S4. Computation time and Memory usage comparison of VAE and EN.** (a) $Log_{10}$ transformed values of computation time in seconds; (b) Memory usage (GB). EN-p.t. (Elastic Net with v100k.P.T variants), VAE: our VAE model using v100k variants, VAE-p.t: VAE-PRS with v100k.P.T variants.

Interaction between 9:273178:G:GA and 3:143021856:G:C (mean_platelet_volume; pAdj = 5.21e-13)

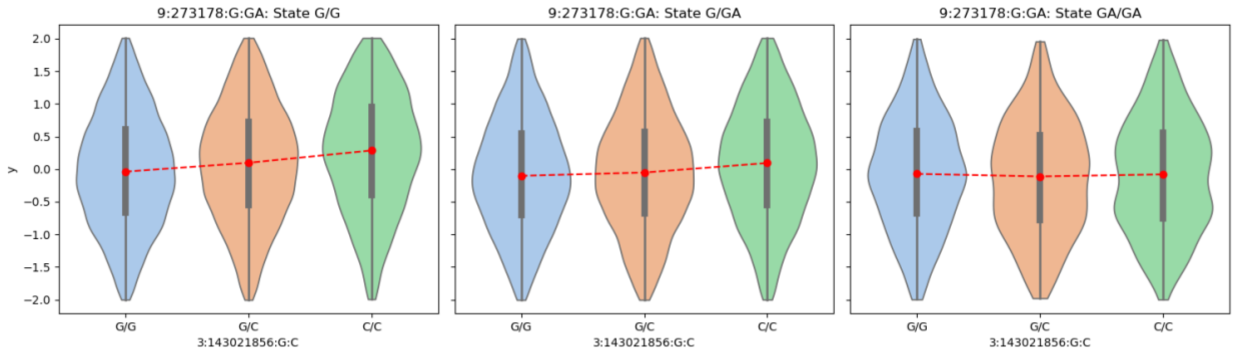


Interaction between 3:143023597:C:T and 3:143021856:G:C (mean_platelet_volume; pAdj = 3.35e-09)

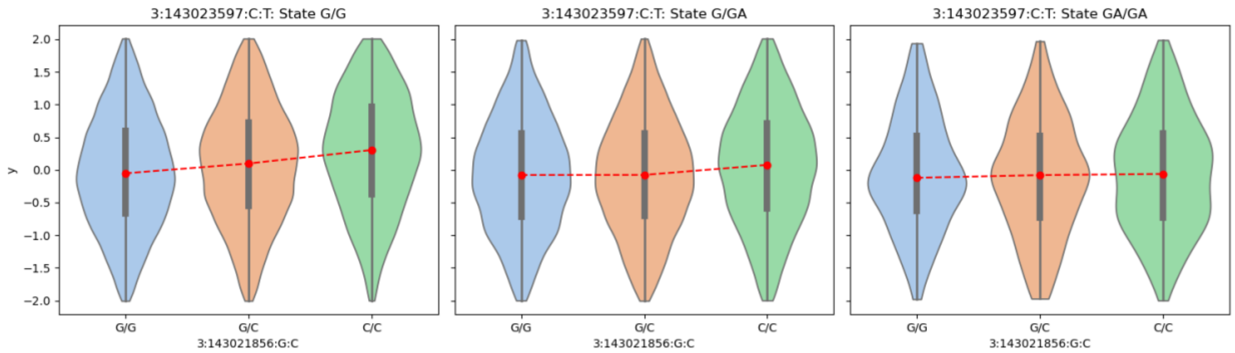


Interaction between 9:273178:G:GA and 3:143023597:C:T (mean_platelet_volume; pAdj = 2.93e-05)

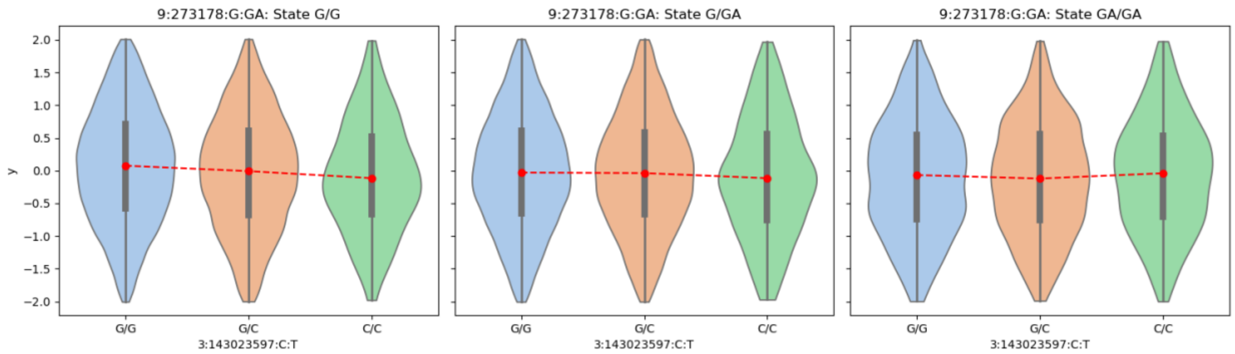

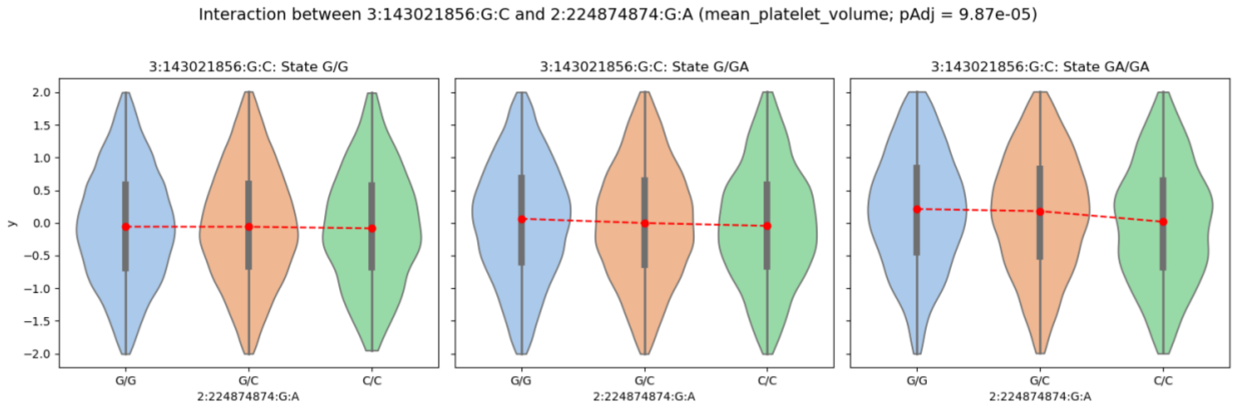


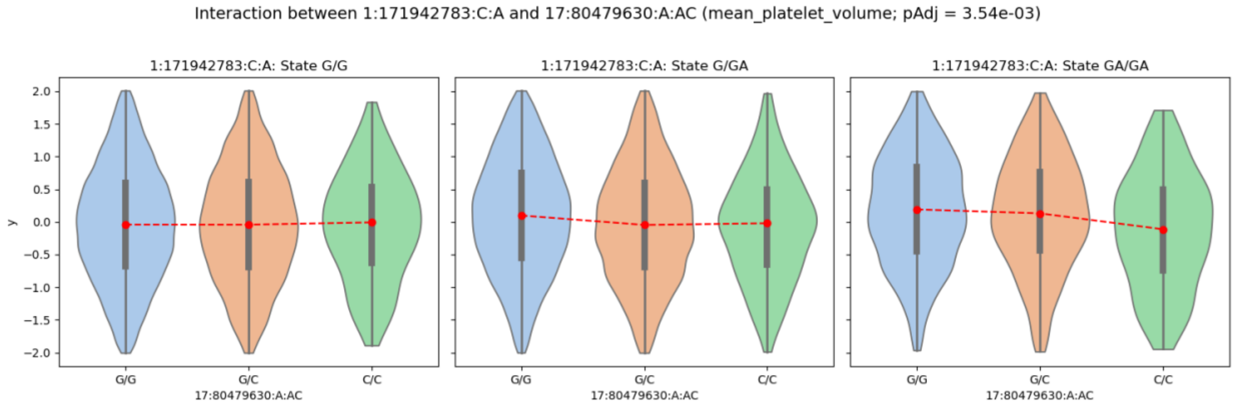


**Supplementary Figure 5. Interaction Analysis for the MPV Trait for top 5 significant pairs of variants.** Each figure illustrates the interaction between two variants, as specified in the title, with the associated p-value of the interaction term also indicated in the title. The change in the slope of the red lines indicates the significance of interaction.

### Supplemental Methods

#### VAE-PRS model framework

Our VAE-PRS for blood cell traits was constructed leveraging a supervised regression model within a Variational Autoencoder (VAE) architecture, as delineated in **Figure 1**. Employing PyTorch Lightning[1], the VAE-PRS method applies an additive encoding (0/1/2) to genotype data before processing through a multilayer perceptron encoder that reduces dimensionality to a latent space. Subsequently, a reparameterization technique is employed to capture the genotype and phenotype distribution characteristics. Finally, the model utilizes a decoder to reconstruct the input from the latent representation, alongside a regression component that predicts continuous phenotypic outcomes.

Mean squared error loss is used as reconstruction loss for both the genotype and the trait of interest. The model is trained to optimize a loss function that balances the reconstruction error of the decoder and the regressor, along with the KL divergence of the latent space distribution. Specifically, our loss function is specified as below

$$\mathcal{L}(\mathbf{X},\mathbf{Y}) = \alpha \cdot \frac{1}{N}\sum_{i=1}^{N}(y_i - \hat{y}_i)^2 + (1-\alpha)\cdot\left(\frac{1}{M \times N}\sum_{i=1}^{M}\sum_{j=1}^{N}(x_{ij} - \hat{x}_{ij})^2 + D_{KL}(\mathbf{q}(\mathbf{z}|\mathbf{X})||\mathbf{p}(\mathbf{z}))\right),$$

where N denotes the number of samples; M denotes the number of features (genetic variants); $y_i$ and $\hat{y}_i$ represent the true and predicted phenotypes, respectively; $x_{ij}$ and $\hat{x}_{ij}$ represent the original and reconstructed genotype values, respectively; $D_{KL}(\mathbf{q}(\mathbf{z}|\mathbf{X})||\mathbf{p}(\mathbf{z}))$ denotes the KL divergence between the approximate posterior distribution $\mathbf{q}(\mathbf{z}|\mathbf{X})$ and the prior distribution, $\mathbf{p}(\mathbf{z})$. The parameter $\alpha$ is the weight for the regression loss, which was treated as a hyperparameter to balance the contribution of each error term to the overall training/validation loss. In our experiments, we chose $\alpha = 0.7$.

When implementing the model in python, all data are loaded with a batch size equal to 64. A learning rate of 1e-6 is used, and an early stop option is included to terminate training and avoid overfitting if the reconstruction loss of phenotype on the validation set has not improved with patient = 8 steps. During training, we recorded both weights and biases to supervise the progress of each error term on both training and validation sets. Experiments are conducted using a single NVIDIA Tesla V100 Volta GPU Accelerator 32GB GPU.

#### Sample split and removal

In this study, we focused on individuals with primarily European ancestry, where ancestry was determined as in our earlier work [2]. The selected samples were partitioned into training, validation, and testing sets according to an 8:1:1 ratio. This stratified random sampling approach ensures that each set is representative of the overall distribution of the data. The training set comprises 80% of the dataset, utilized for model fitting; the validation set accounts for 10%, used for parameter tuning and model selection; and the testing set also represents 10%, reserved for the final evaluation of the model performance. In addition, any samples in the testing set that were related to those in the training/validation set ($r > 0.1$) were removed to avoid potential biases due to relatedness between samples.

#### GWAS

**Genotype data.** We leveraged UKB imputed data[3] with HRC and UK10K as imputation reference panels. First, we converted the imputed data in BGEN file format into PLINK format using PLINK2 software[4]. Plink_pipelines[5] was then used to save the genotype matrices for each individual into individual numpy arrays. These steps transformed the data into a format more readily usable in Python for the VAE-PRS, EN, and MLP models.

**Phenotype data.** We focused on 16 blood cell traits from UK Biobank. Following the same procedure as described before[2,6], we adjusted the covariates with a linear model to account for potential confounding effects, including age, sex, recruitment center, genotyping array, and first 10 PCs. Inverse normalized transformation was applied to ensure a normal distribution. We then performed GWAS with the obtained residuals, which were also used as the input outcome (y in **Figure 1**) in the VAE-PRS model.

**Association.** Genome-wide association study (GWAS) was conducted for PRSice and variant selection in this paper using Regenie v3.1.3[7] with size of the genotype blocks = 1000 and minimum minor allele count (MAC) when testing variants = 20.

#### Other PRS methods

Besides VAE-PRS, we also constructed PRS for blood cell traits using alternative methods for comparison, including C+T method implemented in PRSice2[8], Elastic Net (EN)[9], and multilayer perceptron (MLP).

**PRSice 2 (C+T)**. PRSice2[8] implements a clumping and p-value thresholding technique to determine genetic risk by aggregating the effects of genetic variants identified in genome-wide association studies, with a clumping threshold of $r^2$=0.5 is used. The optimal p-value threshold is tuned based on the testing set with in-sample LD. In this sense, results from PRSice 2 are “overfitted”.

**Elastic Net (EN)**. EN approach, noted for its superior performance in a prior PRS study for blood cell traits[9], employs L1 and L2 regularization penalties to facilitate model sparsity and mitigate overfitting, respectively. Due to the substantial memory and computational demands of EN, it was only applied to variants from conditional analysis and those pruned with LD $r^2$<0.1 and a p-value below 1e-6 in the GWAS summary statistics.

**MLP**. Furthermore, we also designed a three-layer MLP for supervised regression on the traits of interest, utilizing mean squared error for loss calculation and incorporating an early stopping mechanism with a patience of 8 steps to prevent overfitting.

We evaluated PRS using partial $R^2$, which was calculated as the squared Pearson correlation between PRS and phenotype residuals after correcting for covariates and inverse-normal transformation.

#### Assessment of the impact of sample size

The impact of training data volume on deep learning model predictions was examined using VAE-PRS for 16 blood cell phenotypes, with sample sizes from the UK Biobank varied to include 10k, 50k, 100k, 150k, 200k, and 350k individuals. Individuals in each dataset were selected randomly from the full training individuals, followed by a GWAS for each corresponding set. PRS models were constructed as previously described using VAE-PRS, PRSice2 and EN.

**Feature Importance**

The significance of genetic markers and their interactions in the VAE-PRS model was assessed using SHAP (SHapley Additive exPlanations)[10] values for feature importance. SHAP values for each genetic variant were computed using the DeepExplainer module on a cohort of 100 randomly chosen individuals from the test set. These values reflect the mean impact of a variant on the model's predictions, where larger absolute SHAP values denote greater influence, and smaller values indicate lesser importance.

We plotted SHAP feature importance scores against GWAS -log10 p-values and effect sizes for each trait with the mirror plot option from the 'karyoploteR' package in R. Genome build was in hg38.

**Interaction analysis**

To ascertain the contribution of interaction terms to our model's predictive capacity, we ranked variants by their SHAP importance scores, isolating the top 100 for further analysis. Variants demonstrating a Pearson correlation coefficient exceeding 0.9 were excluded to avoid multicollinearity. The selected variants were paired and analyzed for interaction effects using an Ordinary Least Squares (OLS) regression model. Upon adjusting for multiple testings, five variant pairs exhibited significant interaction terms, each with a False Discovery Rate (FDR) below the 0.05 threshold.

**Transfer Learning for non-EUR population**

To investigate the predictive performance of VAE-PRS on non-European population, we fine-tuned on UKB African (AFR) subjects (defined in our previous work[2]) using our VAE-PRS models pre-trained on UKB EUR samples with top 100k variants. A total of 7.8k UKB AFR subjects with blood cell traits were divided into training, validation, and testing sets according to an 8:1:1 ratio. We also directly applied our VAE model pre-trained on UKB EUR samples without fine-tuning as a comparison.

#### References


1. Falcon W, The PyTorch Lightning team. PyTorch Lightning. Published online March 2019. doi:10.5281/zenodo.3828935

2. Sun Q, Graff M, Rowland B, et al. Analyses of biomarker traits in diverse UK biobank participants identify associations missed by European-centric analysis strategies. *J Hum Genet*. 2022;67(2):87-93. doi:10.1038/s10038-021-00968-0

3. Bycroft C, Freeman C, Petkova D, et al. The UK Biobank resource with deep phenotyping and genomic data. *Nature*. 2018;562(7726):203-209. doi:10.1038/s41586-018-0579-z

4. Chang CC, Chow CC, Tellier LC, Vattikuti S, Purcell SM, Lee JJ. Second-generation PLINK: rising to the challenge of larger and richer datasets. *GigaScience*. 2015;4:7. doi:10.1186/s13742-015-0047-8

5. Sigurdsson AI, Westergaard D, Winther O, et al. Deep integrative models for large-scale human genomics. *bioRxiv*. Published online January 1, 2021:2021.06.11.447883. doi:10.1101/2021.06.11.447883

6. Rowland B, Venkatesh S, Tardaguila M, et al. Transcriptome-wide association study in UK Biobank Europeans identifies associations with blood cell traits. *Hum Mol Genet*. 2022;31(14):2333-2347. doi:10.1093/hmg/ddac011

7. Mbatchou J, Barnard L, Backman J, et al. Computationally efficient whole-genome regression for quantitative and binary traits. *Nat Genet*. 2021;53(7):1097-1103. doi:10.1038/s41588-021-00870-7

8. Choi SW, O'Reilly PF. PRSice-2: Polygenic Risk Score software for biobank-scale data. *GigaScience*. 2019;8(7):giz082. doi:10.1093/gigascience/giz082

9. Xu Y, Vuckovic D, Ritchie SC, et al. Machine learning optimized polygenic scores for blood cell traits identify sex-specific trajectories and genetic correlations with disease. *Cell Genomics*. 2022;2(1):100086. doi:10.1016/j.xgen.2021.100086

10. Rodríguez-Pérez R, Bajorath J. Interpretation of machine learning models using shapley values: application to compound potency and multi-target activity predictions. *J Comput Aided Mol Des*. 2020;34(10):1013-1026. doi:10.1007/s10822-020-00314-0

11. Gel B, Serra E. karyoploteR: an R/Bioconductor package to plot customizable genomes displaying arbitrary data. *Bioinforma Oxf Engl*. 2017;33(19):3088-3090. doi:10.1093/bioinformatics/btx346
